# Metabolic network models of *Gardnerella* pangenome identify interactions in the vaginal environment

**DOI:** 10.1101/2022.07.18.500544

**Authors:** Lillian R Dillard, Emma M Glass, Amanda L Lewis, Krystal Thomas-White, Jason A Papin

## Abstract

*Gardnerella* is the primary pathogenic bacterial genus present in the polymicrobial infection known as bacterial vaginosis (BV). Despite BV’s high prevalence and associated chronic and acute women’s health impacts, the *Gardnerella* pangenome is largely uncharacterized at both the genetic and functional metabolic level. Here, we used genome scale metabolic models to characterize *in silico* the *Gardnerella* pangenome metabolic content and assessed metabolic functional capacity within a BV positive cervicovaginal fluid context. Metabolic capacity varied widely across the pangenome, with 38.15% of all reactions as core to the genus, compared to 49.6% of reactions identified as unique to a smaller subset of species. Four genes – *gpsA, fas, suhB, psd* – were identified as core essential genes, critical for *in silico* metabolic function of all analyzed bacterial species in the *Gardnerella* genus. Further understanding of these core essential metabolic functions could inform novel therapeutic strategies to treat BV. These data represent the first metabolic modelling of the *Gardnerella* pangenome and illustrate strain-specific interactions with the vaginal metabolic environment across the pangenome.

## Introduction

Bacterial vaginosis (BV) is one of the most common vaginal conditions in reproductive-age women with vaginal complaints (Schwiertz et al., 2006). BV is a polymicrobial infection of the vagina characterized by low levels of *Lactobacillus*, high levels of diverse anaerobes, a vaginal pH greater than 4.5, thin vaginal discharge, and a fishy odor (Wang, 2000). In North America, BV disproportionately impacts women of color. It is prevalent among Black (33-64%) and Hispanic (31-32%) women compared to their white counterparts (23-35%) (Allsworth et al., 2008; Culhane et al., 2002; Koumans et al., 2007; Muzny and Kardas, 2020; Peebles et al., 2019). The estimated annual healthcare-associated costs for BV treatment globally is $4.8 billion US dollars and an additional $9.6 billion when accounting for BV-associated HIV infection and BV-associated preterm birth(Peebles et al., 2019). The bacterial etiology of BV and mechanisms of pathogenic outcomes remain largely ill-defined. However, since its literature debut in the 1950’s as *‘Haemophilus vaginalis,’ Gardnerella* has consistently been reported as one of the dominant genera in the vagina during BV (Schwebke et al., 2014).

The healthy vagina exhibits normal epithelial and mucosal turnover as a protective mechanism that can help eliminate unwanted colonizers (Amegashie et al., 2017; Muenzner et al., 2010). A vaginal microbiome dominated by *Lactobacillus* is associated with healthy outcomes. This association of *Lactobacillus* healthy outcomes is often attributed to lactic acid production, creating an acidic environment inhospitable to many microorganisms (Tachedjian et al., 2017; Tortelli et al., 2020). Sexual activity, menstruation, hygienic behaviors, hormone fluctuation, antibiotics, and douching can cause changes in the vaginal microbiome and potentially open new niches for pathogens (Gajer et al., 2012; Lopes dos Santos Santiago et al., 2012; Mayer et al., 2015). Despite BV’s pervasiveness and its impact on women’s health, treatment options are limited, and often ineffective. Typical broad spectrum antibiotic courses, specifically metronidazole and clindamycin, present initial success in clearing infection; however, 50% of women experience BV recurrence within twelve months of treatment cessation (Bradshaw and Sobel, 2016). The ineffectiveness of current treatment options highlights the need for a deeper insight into the function of BV-associated organisms such as *Gardnerella*.

There is a significant lack of understanding of the functional metabolism of *Gardnerella*, the bacterial genus that often dominates the vaginal niche in BV. The *Gardnerella* pangenome remains largely uncharacterized, as noted by the rapidly evolving speciation classifications within this genus (Qin and Xiao, 2022). Metabolic predictions using *in silico* analysis offer a unique opportunity to study taxonomic relatedness based on inferred function as opposed to the traditional genetic content-based approach. Previous research has shown that women who are positive for BV typically present with multiple strains of *Gardnerella* (Janulaitiene et al., 2017; Shipitsyna et al., 2019). While multiple, as well as different, strains are present during BV, how these strains differentially interact with vaginal metabolic environment, and the secondary implications for pathogenesis are not well characterized. By understanding these strain level differences, we can begin to define the driving metabolic factors of strain level co-colonization and predict how they interact in BV.

Here, we present the first characterization of *in silico* models reflecting the genetic content and metabolism of the *Gardnerella* pangenome using genome-scale metabolic network reconstructions (GENREs) to allow for the identification of potential antibiotic targets, both strain-specific and conserved, as well as make predictions regarding differential pathogenesis. By defining the conserved features and strain-level variation within the *Gardnerella* pangenome, we can begin to make testable predictions about microbial physiology and give microbial structure to the heterogenous nature of BV.

## Results

### Strain Comparisons

The number of protein coding genes across the 110 *Gardnerella* strains ranged from 434 to 1012, with a median value of 471. The number of genes in the 110 metabolic models ranged from 431 to 688, with a median value of 468. Model metabolites ranged from 782 to 1077, with a median value of 873. Lastly, model reactions ranged from 752 to 1012, with a median of 818. As seen in **Figure 2a**, there are consistently 6 outlier strains across all four categories. A comparison of hierarchical clustering of strains based on full genetic content (including core and peripheral) versus full metabolic content results in an entanglement value of 0.61, which indicates 61% dissimilarity between the two respective dendrograms (**Fig 2b**).

**Figure 1:**
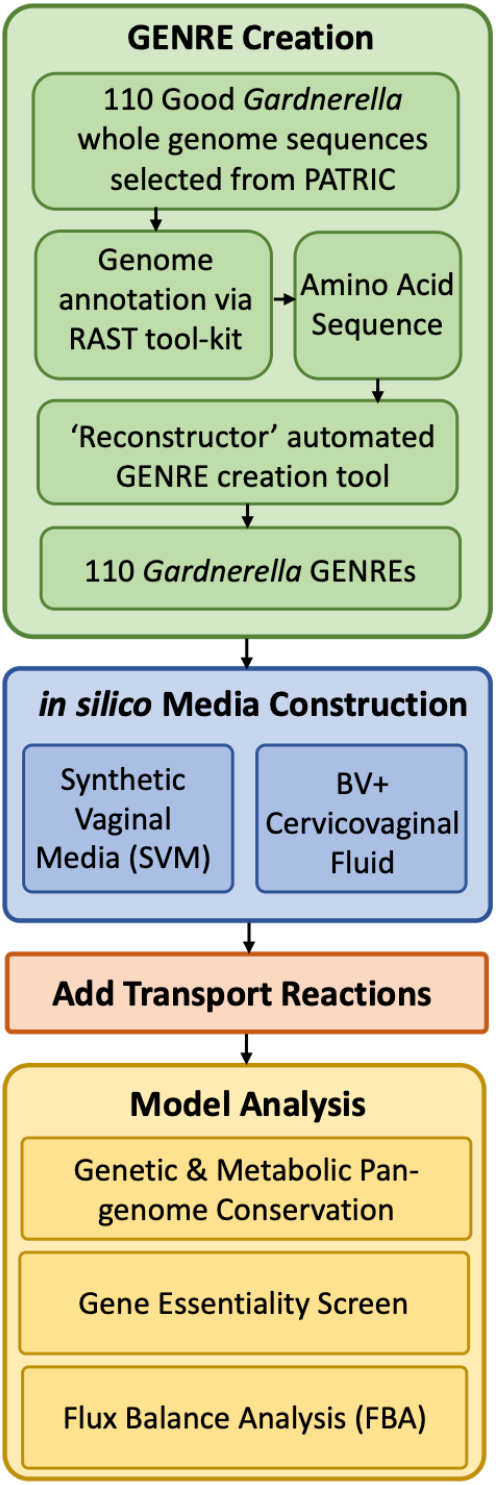
Analysis workflow for pangenome model reconstruction, contextualization, and analysis

**Figure 2:**
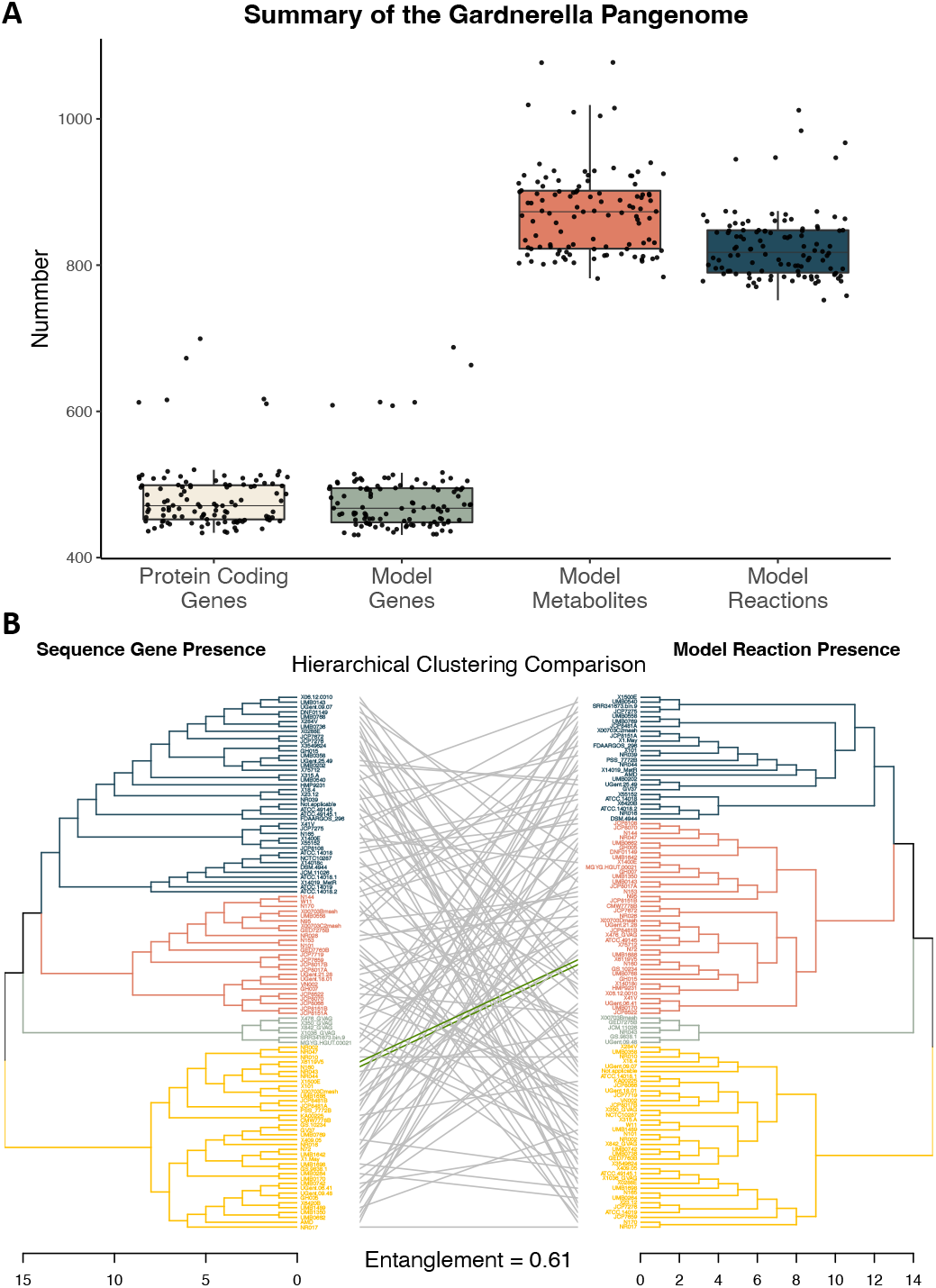
**A)** Summary of pangenome sequence content and corresponding metabolic model content. **B)** Comparison of hierarchical clustering based on protein coding gene presence (left) and model reaction presence (right). Branch color indicates the associated K-means group. Green entanglement lines indicate the strain was placed in the same branch for both dendrograms (note there are only 2 strains that classify in the same groups).

### Genetic and Reaction Conservation

Based on the KEGG ortholog IDs of each gene present across the *Gardnerella* pangenome, 359 genes were considered unique, 90 genes were considered peripheral, and 318 genes were considered core (**Fig 3a**). There is a high degree of conservation of genes implicated in antibiotic resistance across the pangenome. 98% of strains have genes implicated in rifamycin resistance, 88% in mupirocin resistance, 85% in streptogramin resistance, 81% in lincosamide resistance, and 81% in pleuromutilin (**Fig 3b**). In the minority are strains that have genes associated with tetracycline resistance (20% of strains), macrolide resistance (13% of strains), and aminoglycosides resistance (1% of strains).

**Figure 3:**
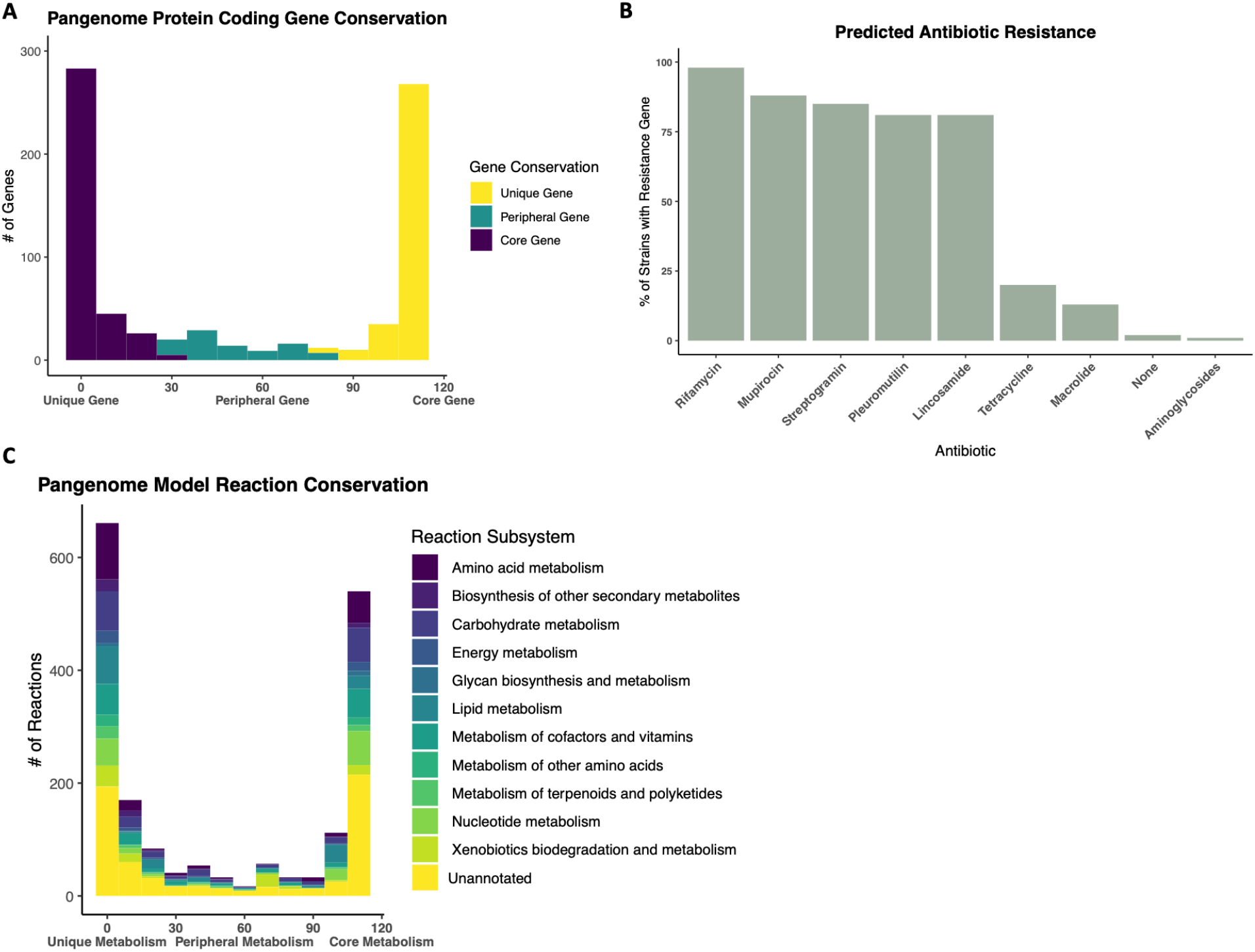
Analysis of pangenome. **A)** Genetic conservation based on protein coding gene presence. **B**) Predicted prevalence of antibiotic resistance genes based on Resistance Gene Identifier protein sequence annotation. **C)** Pangenome metabolic conservation based on model reaction presence.

Based on the 110 metabolic models constructed to represent the *Gardnerella* pangenome, 919 reactions were considered unique, 221 were peripheral reactions, and 695 were core reactions (**Fig 3c**). Of the 209 reactions associated with amino acid metabolism, 122 were unique reactions (58.4%) compared to 70 core reactions (33.5%). Of the 48 reactions associated with terpenoid and polyketide metabolism 29 were unique reactions (60.4%) and only 14 were core reactions (29.2%). Additionally, of the 106 reactions associated with xenobiotics biodegradation and metabolism 55 were unique reactions (51.2%), compared to 22 core reactions (20.8%). Lastly, glycan biosynthesis and metabolism, composed of 18 reactions, was enriched for core reactions (10, 55.6%) compared to unique reactions (6, 33.3%).

### Gene Essentiality

Based on assessed model gene essentiality in both BVCFM and SVM, 57 genes were identified as essential across the pangenome (**Fig 4**). There is near universal essentiality of the four following genes: *gpsA* (K00057), *fas* (K11533), *suhB* (K01092), psd (K01613). KEGG annotations indicate *gpsA* is involved in glycerophospholipid metabolism, *fas* is involved in fatty acid biosynthesis, *suhB* is involved in inositol phosphate metabolism, and *psd* is involved in glycerophospholipid metabolism.

**Figure 4:**
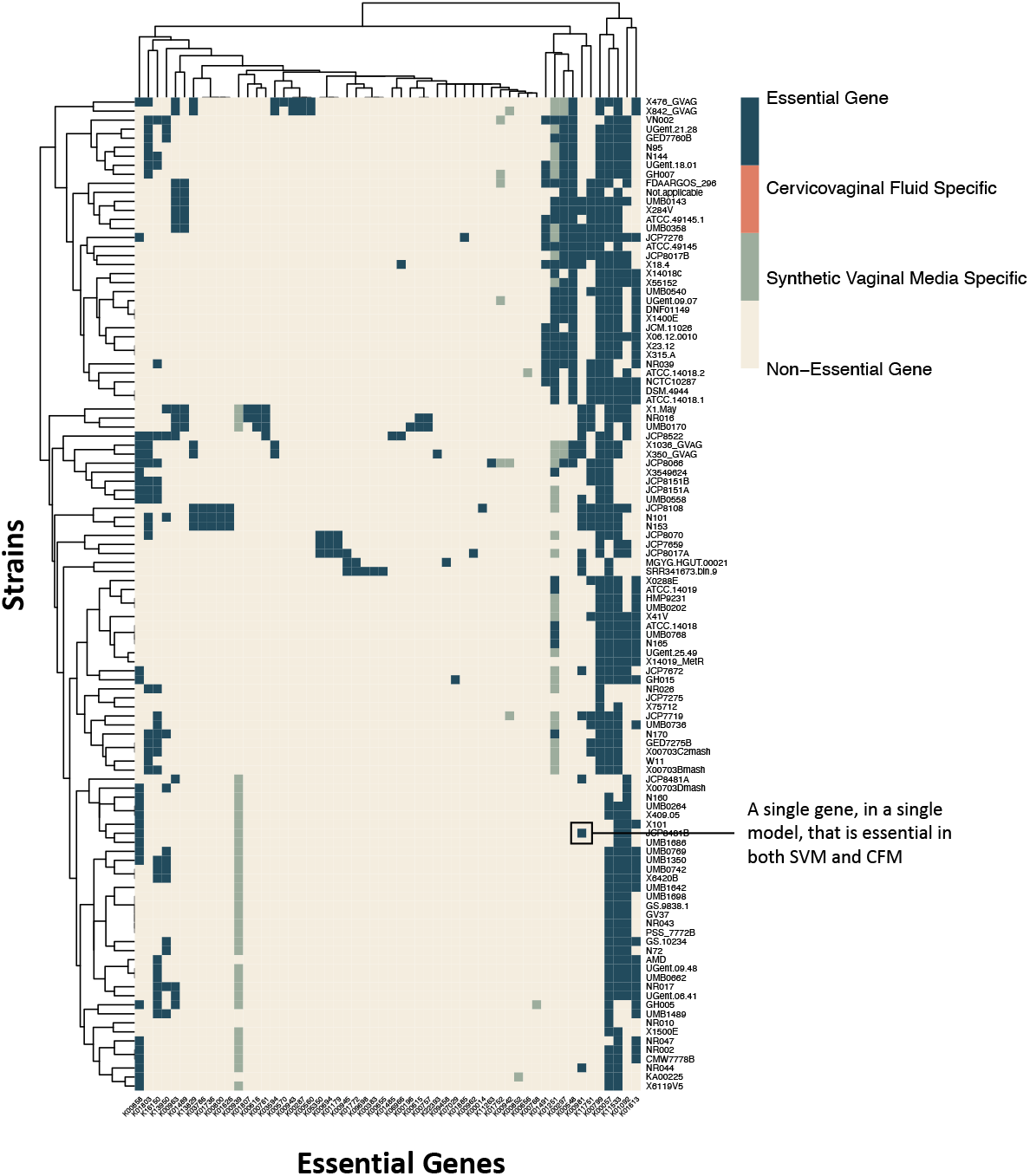
***In silico*** gene essentiality screen across 110 models, contextualized to synthetic vaginal media (SVM) or BV positive cervicovaginal fluid (BVCFM)

### Model Flux Comparisons

When comparing the 110 metabolic models based on flux sampling and dimensionality reduction via tSNE, there is relatively limited clustering based on sample isolation source (**Fig 5**). Samples isolated from the gut and a subset of clinical isolates exhibit clustering. However, when specifically looking at transport flux values alone via tSNE, there is pronounced clustering of gut isolates, blood culture isolates, and more interestingly, the lab strain 14019_MetR isolate (**Fig 6a**). Heatmap comparison of transport reactions with the most varied flux values shows that the subset of models that are exporting L-threonine, chloride, L-valine, and aspartate glutamate also are importing galactose and sodium (**Fig 6b**). Additionally, a small subset of models is significantly importing MAN6P also known as mannose-6-phosphate.

**Figure 5:**
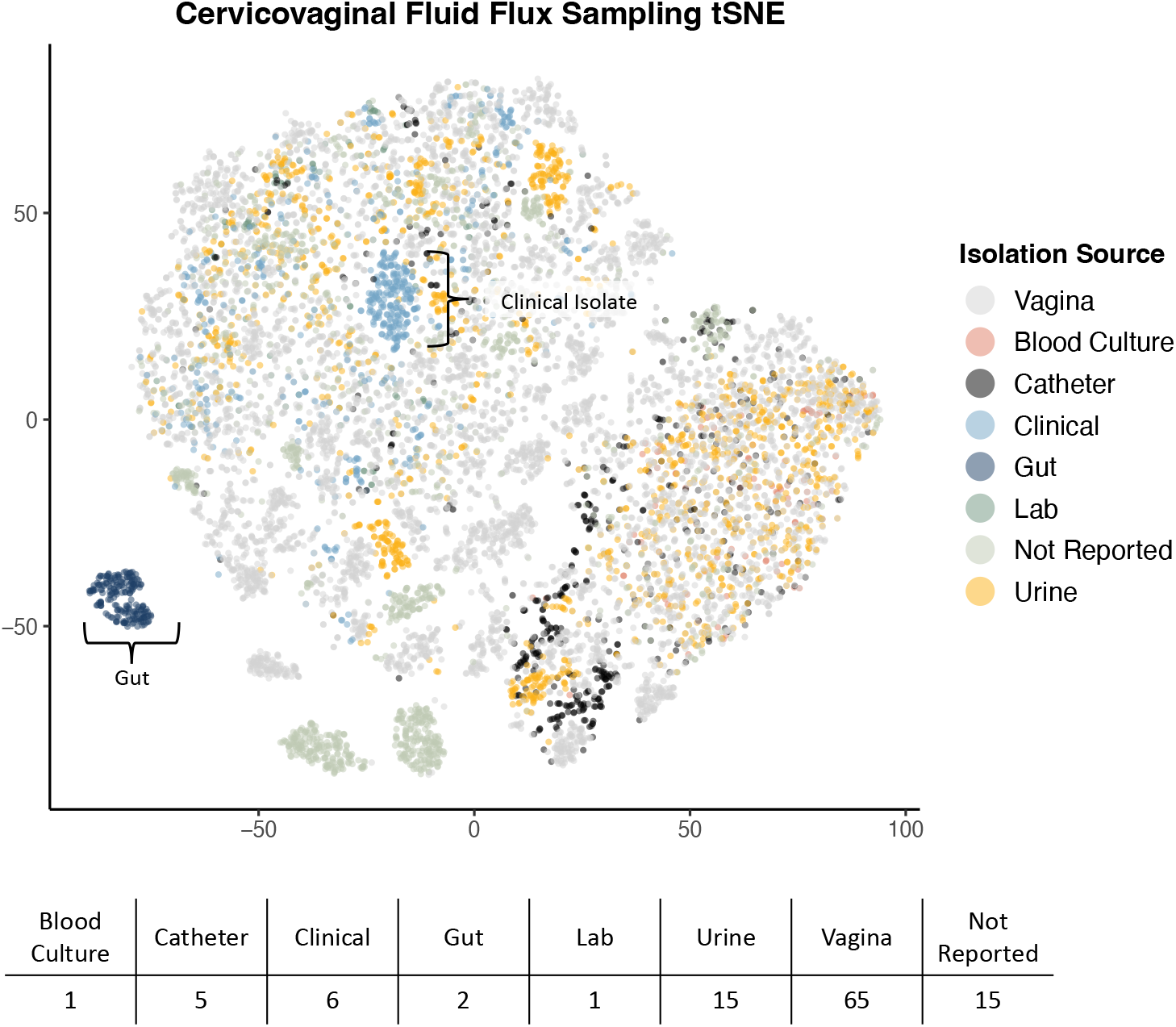
Model flux comparison based on 100 flux samples, across all reactions via tSNE dimensionality reduction and associated number of strains per isolation source.

**Figure 6:**
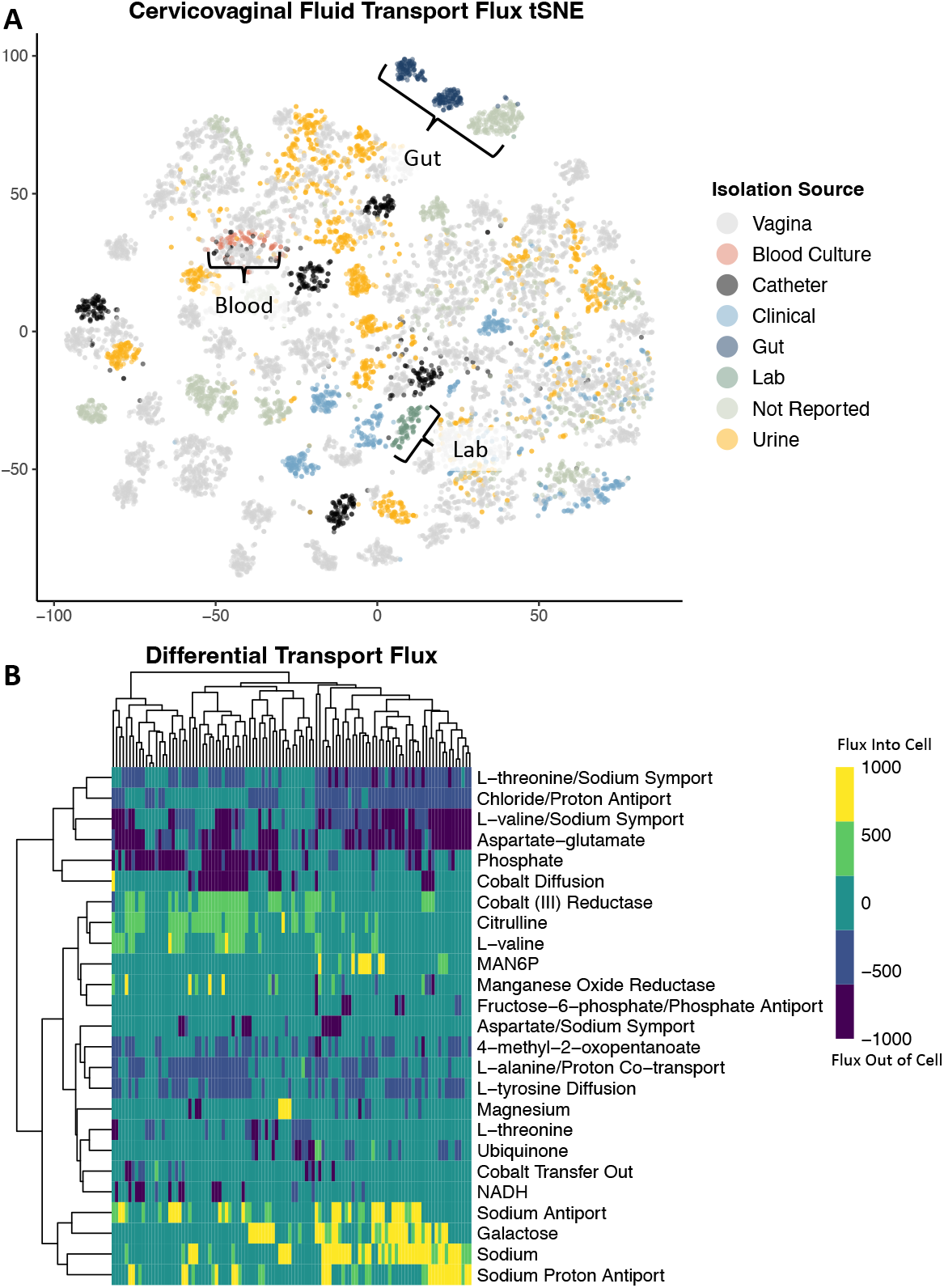
**A)** Model flux comparison based on 100 flux samples, across transport reactions via tSNE dimensionality reduction. **B)** Heatmap of median flux values for each model, based on the top 25 most varied transport flux reaction values. Determined based on 100 flux samples collected via Gapsplit.

## Discussion

The genetic and functional metabolic differences in the *Gardnerella* pangenome remain widely uncharacterized, and *in vitro* analysis of all publicly available strains would require immense time and resources. This lack of information creates a barrier to improving and developing treatment for BV and subsequent reduction of rates of recurrence. Through *in silico* analysis, we have expedited the functional characterization of the *Gardnerella* pangenome. Here, we present the largest set of *Gardnerella* metabolic network reconstructions (GENREs) which characterizes the known *Gardnerella* pangenome. Using these models, we have identified conserved and unique metabolic mechanisms that represent a valuable resource for future treatment development.

### Pangenome Content Comparison

When comparing genetic relatedness to functional relatedness across the *Gardnerella* pangenome, there is a high degree of dissimilarity (61%) between the two dendrograms (**Fig 2b**). This result indicates that genetic content similarity does not directly correlate with metabolic functional similarity when investigating relatedness of *Gardnerella* strains. From an evolutionary perspective, small genetic differences can confer a larger impact on metabolic functionality, as one gene can be involved in multiple reactions. Based on this concept, genetic expression profiles, which may more similarly mirror metabolic functionality (Eisen et al., 1998; Tavazoie et al., 1999; Zaslaver et al., 2004), may offer a more accurate representation of phylogenetic relatedness within the *Gardnerella* pangenome.

In regard to genetic conservation, we specifically investigated the conservation of antibiotic resistance genes due to the translational impact these genes may have on therapeutic approaches for BV infection. We identified which drug classes were associated with the most highly conserved antibiotic resistance genes. These drugs included rifamycin, mupirocin, streptogramin, lincosamide and pleuromutilin (**Fig 3b**). Nitroimidazole class antibiotics, the class that metronidazole falls under, are a part of the resistance gene identifier database, but associated antibiotic resistance genes were not identified within the *Gardnerella* pangenome. Additionally, the large number of antibiotic resistance genes could be a contributing factor to the high rates of BV recurrence, as clindamycin is a class of lincosamide antibiotic (“Lincosamides,” 2016).

The high proportion of unique reactions associated with amino acid metabolism (**Fig 3c**) is supported by previous literature; differential amino acid metabolism can be used to distinguish *Gardnerella* subgroups (Khan et al., 2021). Additionally, the large number of unique reactions associated with xenobiotic biodegradation suggests that at the pangenome level, the *Gardnerella* genus is capable of differentially interacting with pharmaceutical treatments, as well as non-endogenous probiotics (Wang et al., 2019). This result speaks to the need for understanding what *Gardnerella* strains are present in BV to adequately and effectively develop a patient-specific treatment. Interestingly, the large number of unique reactions associated with terpenoid and polyketide metabolism could offer insight as to why some women present with persistent and odorous BV, while others remain asymptomatic. Previous research has shown that polyketide metabolism can function as an antimicrobial agent, allowing the polyketide-producing bacteria to outcompete its microbial competitors (Ridley et al., 2008). In short, this result indicates some *Gardnerella* strains may be uniquely equipped to outcompete others as vaginal microbes. Lastly, glycan-related metabolism is uniquely enriched in the pangenome core metabolism. Vaginal mucus is sialoglycan rich and can be utilized by *Gardnerella* as nutrients if sialidase activity, which hydrolyzes the sialoglycans and frees sialic acid from the glycan chain, is present (Govinden et al., 2018). Interestingly, not all *Gardnerella* strains are sialidase positive (Lewis et al., 2013). Conserved glycan metabolism could indicate inner-pangenome co-evolution of Gardnerella, based on the differences in sialidase activity across strains, as well as potential coevolution with sialic acid catabolizing microbes such as *Fusobacterium* (Agarwal et al., 2020).

### Gene Essentiality

Conserved essential genes could serve as novel targets for drug development. One such essential gene, *gpsA*, is a predicted glycerol-3-phosphate dehydrogenase, and it is involved in phospholipid synthesis. Previous research has identified *gpsA* as a virulence factor in Lyme disease, as well as enhancing nasopharyngeal colonization of *Streptococcus pneumoniae* (Drecktrah et al., 2022; Green et al., 2021). Another conserved essential gene, *suhB*, is an inositol monophosphatase. It has been shown to regulate multiple virulence factors in *Pseudomonas aeruginosa*, as well as play an essential role in *Burkholderia cenocepacia* biofilm formation, motility, and antibiotic resistance (Li et al., 2013; Rosales-Reyes et al., 2012). Based on this previous research, both *gpsA* and *suhB* could be universally essential for driving *Gardnerella* virulence, as well as *Gardnerella* adaptation to the vaginal mucosal environment. *psd* is a phosphatidylserine decarboxylase which plays a role in bacterial membrane biogenesis and has been identified as a potential antimicrobial target (Voelker, 1997). *psd* activity in *Plasmodium falciparum* has been successfully inhibited using 4-quinolinamine compounds (Choi et al., 2016). Due to the essentiality of *psd* in *Gardnerella*, 4-quinolinamine compounds may serve as a starting point for novel BV treatment development. Another conserved essential gene, *fas*, is involved in fatty acid biosynthesis, thus playing an essential role in bacterial membrane construction. Because fatty acid synthase type II (FASII) is bacteria specific, *fas* may be a potential novel BV drug target.

### Flux Analysis

By investigating transport-specific flux values, we can make inferences regarding how strains differentially interact with their vaginal metabolic environment. Dimensionality reduction and visualization via tSNE highlights the differential clustering of lab strain isolates compared to strains collected from body sites (**Fig 6a**). This finding emphasizes the gap in understanding of *Gardnerella* metabolic functionality, due to the primary use of lab strains for *in vitro* experimentation. Secondly, the dispersed nature of vaginal isolates indicates wide variation in functional metabolism of vaginal microbiome strains, and subsequently strains involved in BV (**Fig 6b**). Differential import of galactose could indicate strain-level variation in energy source (Benito et al., 1986). There is a small set of strains that have high levels of mannose-6-phosphate import. The mannose-6-phosphate transport reaction was originally characterized in the *Bacillus subtilis 168* published metabolic model (Oh et al., 2007). *Bacillus subtilis* has been isolated cervicovaginal fluid, and identified via 16S sequencing (das Purkayastha et al., 2019). Mannose-6-phosphate is an essential ligand for the mannose-6-phosphate enzyme, a key enzyme for lysosomal function (Gary-Bobo et al., 2007). Previous research has shown that interrupting mannose-6-phosphate receptor transport from the golgi to the endosome reduces lysosomal function and inhibits host lysosome elimination of infection (Chen et al., 2004; Sun et al., 2021). These findings suggest that some strains of *Gardnerella* may sequester mannose-6-phosphate as a mechanism of evading host lysosomal clearance. This phenotype would result in disordered vaginal epithelial cell function, due to lack of waste removal and would likely be concordant with increased inflammation and oxidative stress (Ferreira and Gahl, 2022; Vuolo et al., 2021).

*Gardnerella* is the one of the most abundant genera present in BV. Despite the high prevalence of BV and the associated negative health impacts, the *Gardnerella* pangenome is largely uncharacterized, at both the genetic and metabolic function level. Using *in silico* analysis via genome-scale metabolic models and vaginal metabolic environment contextualization, we studied 110 *Gardnerella* strains. Through these analyses we discovered that genetic relatedness does not necessarily translate to functional relatedness among *Gardnerella* strains. These findings highlight that BV research should not be overly dependent on genetic relatedness of strains, and instead focus on functional understanding of the *Gardnerella* pangenome, in order to design effective interventions at a strain level. Conserved gene essentiality predictions - specifically *gpsA, fas, suhB, psd* - could inform identification of novel drugs that target this large, diverse genus. While these genes are found in other organisms, such as *Escherichia coli*, there are no current antibiotics available that target any of these four genes. Lastly, *Gardnerella* strains interact differently with their vaginal metabolic environment across the pangenome. This result suggests inner-pangenome co-evolution regarding metabolic niche development, and potential mutualism and commensalism. These discoveries serve as a starting point for developing a deeper understanding of patient-level variation in BV and its impact on health outcomes, infection, and translating these findings to develop personalized medicine therapeutics

## Methods

### Model construction and contextualization

To perform metabolic characterization of the *Gardnerella* pangenome *in silico*, we identified 110 *Gardnerella* whole genome sequences considered “good” quality from the PATRIC 3.6.12 database. PATRIC guidelines define good as “a genome that is sufficiently complete (80%), with sufficiently low contamination (10%)”, and amino acid sequences that are at least 87% consistent with known protein sequence. The corresponding amino acid sequence of these 110 strains were then annotated via RAST 2.0 (Aziz et al., 2008; Brettin et al., 2015; Overbeek et al., 2014). The Reconstructor algorithm was used to create GENREs for each strain (Jenior et al., in preparation). After network construction, two *in silico* media conditions were defined, synthetic vaginal media (SVM) and bacterial vaginosis-positive cervicovaginal fluid media (BVCFM). The SVM condition was based on the composition of previously defined *in vitro* media specific for growth of vaginal microflora (Geshnizgani and Onderdonk, 1992). The BVCFM condition was based on previous metabolomics analysis of cervicovaginal fluid collected from both healthy and BV-positive patients (Vitali et al., n.d.). The BVCFM *in silico* media was enriched for metabolites that were significantly higher in BV-positive cervicovaginal fluid samples in order to specifically investigate metabolic functionality in the disease state. Transport reactions absent from initial model construction but required for *in silico* media metabolite usage were added as needed to the reconstructions, in addition to respective exchange reactions **(Fig 1).**

### Model comparisons

Gene presence for each model was initially determined by the BLASTp output annotations (Madden, 2002). For each BLASTp protein coding gene, the associated KEGG Ortholog was identified and used to construct a binary matrix based on presence or absence of the gene in each model. In parallel, a model reaction presence binary matrix was also constructed. Respective dendrograms of gene presence and reaction presence were constructed using the dendextend package in R (Galili, 2015). Dissimilarity matrices were constructed using the Sørensen–Dice method (Dice, 1945). Hierarchical clustering was calculated using the Ward method (Ward, 1963). Entanglement values of the two dendrograms were calculated using the entanglement function in dendextend. Four k-means clusters are represented by branch coloring for each of the respective dendrograms. For assessing protein coding gene conservation across models, the binary matrix of gene presence was used to determine how many models a gene was or was not present in. Genes present in >75% of models were considered core genes; genes present in 25-75% of models were considered peripheral genes; and genes present in <25% of models were considered unique genes. An analogous metric was used to define model reaction conservation. Reaction subsystems were determined based on the corresponding KEGG reaction metabolic subsystem annotation. Predicted antibiotic resistance was determined with the Resistance Gene Identifier 5.2.1 platform using the amino acid sequence of each strain (Alcock et al., 2020). Genes were considered a match if there was >80% regional match based on protein sequence.

### Gene essentiality

Corresponding exchange reactions were opened (flux bounds of −1000 to 1000) for associated components of the SVM and BVCFM to create two contextualized models for each of the 110 strains. Gene essentiality was determined using the gene essentiality function in the COBRApy toolbox (Ebrahim et al., 2013). This function simulates single gene deletions for every gene present in a model. If a deletion results in >80% reduction in flux through the objective function (biomass synthesis) as determined from simulation of the wildtype condition in the respective media of interest, the gene is categorized as essential. The KEGG ortholog values for each essential gene were identified and used for further analysis. After running the gene essentiality screen, a heatmap was generated using the pheatmap package in R (Kolde, 2019). A Euclidean distance matrix was constructed and then clustered using the complete clustering method.

### Flux comparison

Using the COBRApy-compatible GAPSPLIT function, 100 flux samples were collected for each model in the cervicovaginal media context (Keaty et al., 2020). To better understand inherent clustering of strains based on simulated flux distributions, we utilized dimensionality reduction and visualization via tSNE on the collected flux samples. tSNE analysis was run via the sci-kit learn sklearn.manifold package in python using default parameters (Maaten et al., 2008; Pedregosa et al., 2012). Strain metadata from PATRIC was used to map sample isolation sources for tSNE visualization. Additionally, transport reactions were isolated from the flux sampling data, and the median values for each model’s transport reaction were used to create a heatmap comparing the top 25 most variable transport reaction fluxes across models. The pheatmap library in R was used for heatmap construction, which uses Euclidean distance and complete clustering to determine hierarchical structure.

## Bibliography

Agarwal, K., Robinson, L.S., Aggarwal, S., Foster, L.R., Hernandez-Leyva, A., Lin, H., Tortelli, B.A., O’Brien, V.P., Miller, L., Kau, A.L., Reno, H., Gilbert, N.M., Lewis, W.G., Lewis, A.L., 2020. Glycan cross-feeding supports mutualism between Fusobacterium and the vaginal microbiota. PLoS Biology 18. https://doi.org/10.1371/JOURNAL.PBIO.3000788

Alcock, B.P., Raphenya, A.R., Lau, T.T.Y., Tsang, K.K., Bouchard, M., Edalatmand, A., Huynh, W., Nguyen, A.L. v., Cheng, A.A., Liu, S., Min, S.Y., Miroshnichenko, A., Tran, H.K., Werfalli, R.E., Nasir, J.A., Oloni, M., Speicher, D.J., Florescu, A., Singh, B., Faltyn, M., Hernandez-Koutoucheva, A., Sharma, A.N., Bordeleau, E., Pawlowski, A.C., Zubyk, H.L., Dooley, D., Griffiths, E., Maguire, F., Winsor, G.L., Beiko, R.G., Brinkman, F.S.L., Hsiao, W.W.L., Domselaar, G. v., McArthur, A.G., 2020. CARD 2020: antibiotic resistome surveillance with the comprehensive antibiotic resistance database. Nucleic Acids Res 48, D517–D525. https://doi.org/10.1093/NAR/GKZ935

Allsworth, J.E., Lewis, V.A., Peipert, J.F., 2008. Viral sexually transmitted infections and bacterial vaginosis: 2001-2004 National Health and Nutrition Examination Survey data. Sex Transm Dis 35, 791–796. https://doi.org/10.1097/OLQ.0B013E3181788301

Amegashie, C.P., Gilbert, N.M., Peipert, J.F., Allsworth, J.E., Lewis, W.G., Lewis, A.L., 2017. Relationship between nugent score and vaginal epithelial exfoliation. PLoS One 12. https://doi.org/10.1371/JOURNAL.PONE.0177797

Aziz, R.K., Bartels, D., Best, A., DeJongh, M., Disz, T., Edwards, R.A., Formsma, K., Gerdes, S., Glass, E.M., Kubal, M., Meyer, F., Olsen, G.J., Olson, R., Osterman, A.L., Overbeek, R.A., McNeil, L.K., Paarmann, D., Paczian, T., Parrello, B., Pusch, G.D., Reich, C., Stevens, R., Vassieva, O., Vonstein, V., Wilke, A., Zagnitko, O., 2008. The RAST Server: rapid annotations using subsystems technology. BMC Genomics 9. https://doi.org/10.1186/1471-2164-9-75

Benito, R., Vazquez, J.A., Berron, S., Fenoll, A., Saez-Neito, J.A., 1986. A modified scheme for biotyping Gardnerella vaginalis. Journal of Medical Microbiology 21, 357–359. https://doi.org/10.1099/00222615-21-4-357/CITE/REFWORKS

Bradshaw, C.S., Sobel, J.D., 2016. Current Treatment of Bacterial Vaginosis-Limitations and Need for Innovation. Journal of Infectious Diseases 214, S14–S20. https://doi.org/10.1093/infdis/jiw159

Brettin, T., Davis, J.J., Disz, T., Edwards, R.A., Gerdes, S., Olsen, G.J., Olson, R., Overbeek, R., Parrello, B., Pusch, G.D., Shukla, M., Thomason, J.A., Stevens, R., Vonstein, V., Wattam, A.R., Xia, F., 2015. RASTtk: a modular and extensible implementation of the RAST algorithm for building custom annotation pipelines and annotating batches of genomes. Sci Rep 5. https://doi.org/10.1038/SREP08365

Chen, J.J., Zhu, Z., Gershon, A.A., Gershon, M.D., 2004. Mannose 6-phosphate receptor dependence of varicella zoster virus infection in vitro and in the epidermis during varicella and zoster. Cell 119, 915–926. https://doi.org/10.1016/J.CELL.2004.11.007

Choi, J.Y., Kumar, V., Pachikara, N., Garg, A., Lawres, L., Toh, J.Y., Voelker, D.R., ben Mamoun, C., 2016. Characterization of Plasmodium phosphatidylserine decarboxylase expressed in yeast and application for inhibitor screening. Mol Microbiol 99, 999. https://doi.org/10.1111/MMI.13280

Culhane, J.F., Rauh, V., McCollum, K.F., Elo, I.T., Hogan, V., 2002. Exposure to chronic stress and ethnic differences in rates of bacterial vaginosis among pregnant women. Am J Obstet Gynecol 187, 1272–1276. https://doi.org/10.1067/MOB.2002.127311

das Purkayastha, S., Bhattacharya, M.K., Prasad, H.K., Upadhyaya, H., Lala, S. das, Pal, K., Das, M., Sharma, G.D., Bhattacharjee, M.J., 2019. Contrasting diversity of vaginal lactobacilli among the females of Northeast India. BMC Microbiology 19. https://doi.org/10.1186/S12866-019-1568-6

Dice, L.R., 1945. Measures of the Amount of Ecologic Association Between Species. Ecology 26, 297–302. https://doi.org/10.2307/1932409

Drecktrah, D., Hall, L.S., Crouse, B., Schwarz, B., Richards, C., Bohrnsen, E., Wulf, M., Long, B., Bailey, J., Gherardini, F., Bosio, C.M., Lybecker, M.C., Samuels, D.S., 2022. The glycerol-3-phosphate dehydrogenases GpsA and GlpD constitute the oxidoreductive metabolic linchpin for Lyme disease spirochete host infectivity and persistence in the tick. PLoS Pathog 18. https://doi.org/10.1371/JOURNAL.PPAT.1010385

Ebrahim, A., Lerman, J.A., Palsson, B.O., Hyduke, D.R., 2013. COBRApy: COnstraints-Based Reconstruction and Analysis for Python. BMC Syst Biol 7. https://doi.org/10.1186/1752-0509-7-74

Eisen, M.B., Spellman, P.T., Brown, P.O., Botstein, D., 1998. Cluster analysis and display of genome-wide expression patterns. Proc Natl Acad Sci U S A 95, 14863–14868. https://doi.org/10.1073/PNAS.95.25.14863

Ferreira, C.R., Gahl, W.A., 2022. Lysosomal Storage Disease. Metabolic Diseases: Foundations of Clinical Management, Genetics, and Pathology 367–440. https://doi.org/10.3233/978-1-61499-718-4-367

Gajer, P., Brotman, R.M., Bai, G., Sakamoto, J., Schütte, U.M.E., Zhong, X., Koenig, S.S.K., Fu, L., Ma, Z., Zhou, X., Abdo, Z., Forney, L.J., Ravel, J., 2012. Temporal dynamics of the human vaginal microbiota. Sci Transl Med 4. https://doi.org/10.1126/SCITRANSLMED.3003605

Galili, T., 2015. dendextend: an R package for visualizing, adjusting and comparing trees of hierarchical clustering. Bioinformatics 31, 3718–3720. https://doi.org/10.1093/BIOINFORMATICS/BTV428

Gary-Bobo, M., Nirdé, P., Jeanjean, A., Morère, A., Garcia, M., 2007. Mannose 6-phosphate receptor targeting and its applications in human diseases. Current Medicinal Chemistry 14, 2945. https://doi.org/10.2174/092986707782794005

Geshnizgani, A.M., Onderdonk, A.B., 1992. Defined medium simulating genital tract secretions for growth of vaginal microflora. Journal of Clinical Microbiology. https://doi.org/10.1128/jcm.30.5.1323-1326.1992

Govinden, G., Parker, J.L., Naylor, K.L., Frey, A.M., Anumba, D.O.C., Stafford, G.P., 2018. Inhibition of sialidase activity and cellular invasion by the bacterial vaginosis pathogen Gardnerella vaginalis. Archives of Microbiology 200, 1129. https://doi.org/10.1007/S00203-018-1520-4

Green, A.E., Howarth, D., Chaguza, C., Echlin, H., Langendonk, R.F., Munro, C., Barton, T.E., Hinton, J.C.D., Bentley, S.D., Rosch, J.W., Neill, D.R., 2021. Pneumococcal Colonization and Virulence Factors Identified Via Experimental Evolution in Infection Models. Molecular Biology and Evolution 38, 2209. https://doi.org/10.1093/MOLBEV/MSAB018

Janulaitiene, M., Paliulyte, V., Grinceviciene, S., Zakareviciene, J., Vladisauskiene, A., Marcinkute, A., Pleckaityte, M., 2017. Prevalence and distribution of Gardnerella vaginalis subgroups in women with and without bacterial vaginosis. BMC Infectious Diseases 17, 1–9. https://doi.org/10.1186/S12879-017-2501-Y/TABLES/1

Jenior ML, Glass EM, Papin JA (in preparation) Reconstructor: A COBRApy compatible tool for automated genome scale metabolic network reconstruction through parsimonious flux-based gapfilling

Keaty, T.C., Keaty, T.C., Jensen, P.A., Jensen, P.A., Jensen, P.A., 2020. Gapsplit: efficient random sampling for non-convex constraint-based models. Bioinformatics 36, 2623–2625. https://doi.org/10.1093/BIOINFORMATICS/BTZ971

Khan, S., Vancuren, S.J., Hill, J.E., 2021. A Generalist Lifestyle Allows Rare Gardnerella spp. to Persist at Low Levels in the Vaginal Microbiome. Microb Ecol 82, 1048–1060. https://doi.org/10.1007/S00248-020-01643-1

Kolde, R., 2019. Package “pheatmap”: Pretty heatmaps. Version 1.0.12 1–8.

Koumans, E.H., Sternberg, M., Bruce, C., McQuillan, G., Kendrick, J., Sutton, M., Markowitz, L.E., 2007. The prevalence of bacterial vaginosis in the United States, 2001-2004; associations with symptoms, sexual behaviors, and reproductive health. Sex Transm Dis 34, 864–869. https://doi.org/10.1097/OLQ.0B013E318074E565

Lewis, N.E., Hixson, K.K., Conrad, T.M., Lerman, J.A., Charusanti, P., Polpitiya, A.D., Adkins, J.N., Schramm, G., Purvine, S.O., Lopez-Ferrer, D., Weitz, K.K., Eils, R., König, R., Smith, R.D., Palsson, B., 2010. Omic data from evolved E. coli are consistent with computed optimal growth from genome-scale models. Mol Syst Biol 6. https://doi.org/10.1038/MSB.2010.47

Lewis, W.G., Robinson, L.S., Gilbert, N.M., Perry, J.C., Lewis, A.L., 2013. Degradation, Foraging, and Depletion of Mucus Sialoglycans by the Vagina-adapted Actinobacterium Gardnerella vaginalis. The Journal of Biological Chemistry 288, 12067. https://doi.org/10.1074/JBC.M113.453654

Li, K., Xu, C., Jin, Y., Sun, Z., Liu, C., Shi, J., Chen, G., Chen, R., Jin, S., Wu, W., 2013. SuhB is a regulator of multiple virulence genes and essential for pathogenesis of Pseudomonas aeruginosa. mBio 4. https://doi.org/10.1128/MBIO.00419-13

Lincosamides, 2016. Meyler’s Side Effects of Drugs 581–588. https://doi.org/10.1016/B978-0-444-53717-1.00980-X

Lopes dos Santos Santiago, G., Tency, I., Verstraelen, H., Verhelst, R., Trog, M., Temmerman, M., Vancoillie, L., Decat, E., Cools, P., Vaneechoutte, M., 2012. Longitudinal qPCR study of the dynamics of L. crispatus, L. iners, A. vaginae, (sialidase positive) G. vaginalis, and P. bivia in the vagina. PLoS One 7. https://doi.org/10.1371/JOURNAL.PONE.0045281

Maaten, L. van der, research, G.H.-J. of machine learning, 2008, undefined, 2008. Visualizing data using t-SNE. jmlr.org 9, 2579–2605.

Madden, T., 2002. The BLAST Sequence Analysis Tool [WWW Document]. The NCBI Handbook. URL https://www.ncbi.nlm.nih.gov/books/NBK21097/ (accessed 6.6.22).

Mayer, B.T., Srinivasan, S., Fiedler, T.L., Marrazzo, J.M., Fredricks, D.N., Schiffer, J.T., 2015. Rapid and Profound Shifts in the Vaginal Microbiota Following Antibiotic Treatment for Bacterial Vaginosis. J Infect Dis 212, 793–802. https://doi.org/10.1093/INFDIS/JIV079

Muenzner, P., Bachmann, V., Zimmermann, W., Hentschel, J., Hauck, C.R., 2010. Human-restricted bacterial pathogens block shedding of epithelial cells by stimulating integrin activation. Science 329, 1197–1201. https://doi.org/10.1126/SCIENCE.1190892

Muzny, C.A., Kardas, P., 2020. A Narrative Review of Current Challenges in the Diagnosis and Management of Bacterial Vaginosis. Sexually Transmitted Diseases 47, 441–446. https://doi.org/10.1097/OLQ.0000000000001178

Oh, Y.K., Palsson, B.O., Park, S.M., Schilling, C.H., Mahadevan, R., 2007. Genome-scale reconstruction of metabolic network in Bacillus subtilis based on high-throughput phenotyping and gene essentiality data. J Biol Chem 282, 28791–28799. https://doi.org/10.1074/JBC.M703759200

Overbeek, R., Olson, R., Pusch, G.D., Olsen, G.J., Davis, J.J., Disz, T., Edwards, R.A., Gerdes, S., Parrello, B., Shukla, M., Vonstein, V., Wattam, A.R., Xia, F., Stevens, R., 2014. The SEED and the Rapid Annotation of microbial genomes using Subsystems Technology (RAST). Nucleic Acids Res 42. https://doi.org/10.1093/NAR/GKT1226

Pedregosa, F., Varoquaux, G., Gramfort, A., Michel, V., Thirion, B., Grisel, O., Blondel, M., Müller, A., Nothman, J., Louppe, G., Prettenhofer, P., Weiss, R., Dubourg, V., Vanderplas, J., Passos, A., Cournapeau, D., Brucher, M., Perrot, M., Duchesnay, É., 2012. Scikit-learn: Machine Learning in Python.

Peebles, K., Velloza, J., Balkus, J.E., McClelland, R.S., Barnabas, R. v., 2019. High Global Burden and Costs of Bacterial Vaginosis: A Systematic Review and Meta-Analysis. Sexually Transmitted Diseases 46, 304–311. https://doi.org/10.1097/OLQ.0000000000000972

Qin, H., Xiao, B., 2022. Research Progress on the Correlation Between Gardnerella Typing and Bacterial Vaginosis. Frontiers in Cellular and Infection Microbiology 12. https://doi.org/10.3389/FCIMB.2022.858155

Ridley, C.P., Ho, Y.L., Khosla, C., 2008. Evolution of polyketide synthases in bacteria. Proc Natl Acad Sci U S A 105, 4595–4600. https://doi.org/10.1073/PNAS.0710107105/SUPPL_FILE/10107FIG9.PDF

Rosales-Reyes, R., Saldías, M.S., Aubert, D.F., El-Halfawy, O.M., Valvano, M.A., 2012. The suhB gene of Burkholderia cenocepacia is required for protein secretion, biofilm formation, motility and polymyxin B resistance. Microbiology (Reading) 158, 2315–2324. https://doi.org/10.1099/MIC.0.060988-0

Schwebke, J.R., Muzny, C.A., Josey, W.E., 2014. Role of Gardnerella vaginalis in the pathogenesis of bacterial vaginosis: A conceptual model. Journal of Infectious Diseases 210, 338–343. https://doi.org/10.1093/infdis/jiu089

Schwiertz, A., Taras, D., Rusch, K., Rusch, V., 2006. Throwing the dice for the diagnosis of vaginal complaints? Ann Clin Microbiol Antimicrob 5. https://doi.org/10.1186/1476-0711-5-4

Shipitsyna, E., Krysanova, A., Khayrullina, G., Shalepo, K., Savicheva, A., Guschin, A., Unemo, M., 2019. Quantitation of all Four Gardnerella vaginalis Clades Detects Abnormal Vaginal Microbiota Characteristic of Bacterial Vaginosis More Accurately than Putative G. vaginalis Sialidase A Gene Count. Molecular Diagnosis & Therapy 23, 139. https://doi.org/10.1007/S40291-019-00382-5

Sun, J., Wang, X., Lin, H., Wan, L., Chen, J., Yang, X., Li, D., Zhang, Y., He, X., Wang, B., Dong, M., Zhong, H., Wei, C., 2021. Shigella escapes lysosomal degradation through inactivation of Rab31 by IpaH4.5. J Med Microbiol 70. https://doi.org/10.1099/JMM.0.001382

Tachedjian, G., Aldunate, M., Bradshaw, C.S., Cone, R.A., 2017. The role of lactic acid production by probiotic Lactobacillus species in vaginal health. Res Microbiol 168, 782–792. https://doi.org/10.1016/J.RESMIC.2017.04.001

Tavazoie, S., Hughes, J.D., Campbell, M.J., Cho, R.J., Church, G.M., 1999. Systematic determination of genetic network architecture. Nature Genetics 1999 22:3 22, 281–285. https://doi.org/10.1038/10343

Tian, M., Reed, J.L., 2018. Integrating proteomic or transcriptomic data into metabolic models using linear bound flux balance analysis. Bioinformatics 34, 3882. https://doi.org/10.1093/BIOINFORMATICS/BTY445

Tortelli, B.A., Lewis, W.G., Allsworth, J.E., Member-Meneh, N., Foster, L.R., Reno, H.E., Peipert, J.F., Fay, J.C., Lewis, A.L., 2020. Associations between the vaginal microbiome and Candida colonization in women of reproductive age. Am J Obstet Gynecol 222, 471.e1–471.e9. https://doi.org/10.1016/J.AJOG.2019.10.008

Vitali, B., Cruciani, & F., Picone, G., Parolin, & C., Donders, & G., Laghi, & L., n.d. Vaginal microbiome and metabolome highlight specific signatures of bacterial vaginosis. https://doi.org/10.1007/s10096-015-2490-y

Voelker, D.R., 1997. Phosphatidylserine decarboxylase. Biochim Biophys Acta 1348, 236–244. https://doi.org/10.1016/S0005-2760(97)00101-X

Vuolo, D., do Nascimento, C.C., D’Almeida, V., 2021. Reproduction in Animal Models of Lysosomal Storage Diseases: A Scoping Review. Frontiers in Molecular Biosciences 8, 1113. https://doi.org/10.3389/FMOLB.2021.773384/BIBTEX

Wang, J., 2000. Bacterial vaginosis. Primary Care Update for OB/GYNS 7, 181–185. https://doi.org/10.1016/S1068-607X(00)00043-3

Wang, Z., He, Y., Zheng, Y., 2019. Probiotics for the Treatment of Bacterial Vaginosis: A Meta-Analysis. Int J Environ Res Public Health 16. https://doi.org/10.3390/IJERPH16203859

Ward, J.H., 1963. Hierarchical Grouping to Optimize an Objective Function. J Am Stat Assoc 58, 236–244. https://doi.org/10.1080/01621459.1963.10500845

Zaslaver, A., Mayo, A.E., Rosenberg, R., Bashkin, P., Sberro, H., Tsalyuk, M., Surette, M.G., Alon, U., 2004. Just-in-time transcription program in metabolic pathways. Nature Genetics 2004 36:5 36, 486–491. https://doi.org/10.1038/ng1348

